# wavess 1.2: Presenting an HLA-aware within-host virus sequence simulation framework

**DOI:** 10.64898/2026.02.19.706869

**Authors:** Zena Lapp, Thomas Leitner

## Abstract

**Motivation:** Understanding how virus sequences are shaped by selection can inform vaccine design and transmission inference. Modeling within-host evolution to interrogate these questions requires a detailed mechanistic framework that accurately captures sequence diversification. The CD8^+^ cytotoxic T-lymphocyte (CTL) response plays an important role in immune-mediated selection and can leave strong signatures in virus sequences; however, existing sequence-based within-host virus modeling frameworks do not explicitly include a human leukocyte antigen (HLA)-aware CTL response.

**Results:** We extended our previously published within-host sequence evolution simulator, `wavess`, to include an explicit CTL response, and share a method for identifying HLA-specific CTL epitopes given a founder virus sequence. We also updated the model to permit a variable recombination rate, which allows for modeling non-adjacent genes, segmented genomes, and recombination hotspots. These extensions to `wavess` allow for more accurate simulation of viruses and virus genes, particularly in regions of the genome where the immune response is dominated by CTLs (rather than antibodies). It also provides the foundation for investigations of how these newly-added biological mechanisms influence within-host evolution.

**Availability and implementation:** The core of `wavess` is written in Python 3, with helper functions written in R. It is available at https://github.com/MolEvolEpid/wavess.

## Introduction

Forward-in-time simulation of virus sequence evolution in a host can provide fundamental knowledge relevant to public health. It can help tease apart how different selective pressures influence virus survival, which may inform vaccine development. It can also improve our understanding of how to correctly infer transmission networks from sequences under strong selective pressures, which may inform inference tools used to suggest interventions. To investigate these questions, the model must accurately simulate mutational signatures in virus sequences.

An important selective pressure that shapes the virus fitness landscape, and thus sequence signatures, is the CD8^+^ cytotoxic T-lymphocyte (CTL) immune response (Carlson et al., 2015). CTLs recognize non-self peptides presented by human leukocyte antigen (HLA) class I molecules and mount an immune response against infected cells. There is strong pressure on the virus to escape this response, which it can do by mutating such that the peptide can no longer be presented by the host’s HLA molecules. This escape signature can be identified in virus sequence data from infected individuals (Fischer et al., 2010). However, sequence-based within-host simulators (Arenas, 2013; Jariani et al., 2019; Perera et al., 2025) do not explicitly model a CTL immune response that can vary depending on the host’s HLA molecules, meaning that the timing of escape from the CTL response and genetic signatures of escape cannot be explored.

We previously developed a model and R package, `wavess` (Sambaturu et al., 2025), that simulates within-host virus evolution with recombination and latency under selective pressures including conserved sites, replicative fitness, and a generic adaptive immune response with cross-reactivity. Here, we describe how we updated this model to allow for an explicit CTL immune response. We also briefly describe the addition of variable recombination rates across the sequence, which, while not novel (Arenas, 2013), expands the use cases of the simulator. These features are included in `wavess` version 1.2.

## Model updates

`wavess` is an individual-based forward-simulation model that explicitly simulates virus sequences including mutation, recombination, and selection (Sambaturu et al., 2025). Below, we describe two new `wavess` model components: a host CTL immune response and a variable recombination rate.

### HLA-specific CTL immune response

`wavess` 1.0 modeled a generic adaptive immune response with cross-reactivity and no option for complete escape; however, the CTL response leads to distinctly observable escape mutations, particularly early in infection (Fischer et al., 2010). As immune-driven evolution in many virus genes is mostly due to the host CTL response rather than the antibody response, we modified `wavess` to separately model these two components of the immune system. The B-cell antibody immune response is modeled identically to the original `wavess` immune response. In addition, we added functionality to model complete escape from a CTL response at user-defined amino acid positions. One amino acid is recognized by the immune system; mutation to any other amino acid confers complete and immediate escape from recognition. All T-cell epitopes have an identical user-defined maximum fitness cost *c*_*max*_ (default = 0.5), and each T-cell epitope *i* has a(n optionally) distinct time to reach maximum fitness cost 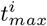, so T-cell responses can mature at different rates. All responses mature linearly from 0 to 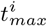 and once 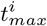 is reached, that cost is maintained for the remainder of the infection. So the fitness of a recognized epitope *i* in generation *t* is defined as:

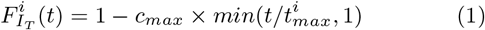

Epitopes that have escaped recognition have a fitness of 1. The T-cell immune fitness for a virus at generation *t* is the product of the individual fitness for each epitope:

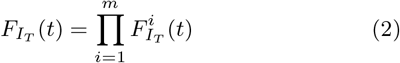

where *m* is the total number of CTL epitopes in the individual.

The overall fitness for a virus at time *t* is the product of the virus’s conserved fitness *F*_*C*_, replicative fitness *F*_*R*_, B-cell (antibody) fitness 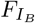, and T-cell (CTL) fitness 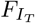:

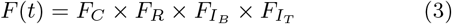

### Variable recombination rate

`wavess` 1.0 used a recombination rate (a combined rate of dual infection followed by recombination) that was identical for all breakpoints. We have since implemented the option to have different rates for each breakpoint between nucleotides to primarily enable computationally efficient modeling of non-adjacent genes in a sequence (without having to model the intervening sequence), but also segmented viruses and recombination hotspots. We convert each user-defined rate into the probability of an odd number of recombination events occurring at a given breakpoint as (1−*e*^−2*nλ*^)*/*2 (which assumes a Poisson process), because if an even number occurs then it is not observable in the sequence. Rates over 3 recombinations per breakpoint per generation lead to a probability of about 0.5. We used GitHub Co-Pilot to help with code optimization.

### Usage updates and availability

We have updated the `wavess` R package vignettes to describe, and provide examples of, the new model options. All updates can be found at https://github.com/MolEvolEpid/wavess.

## Methods

We applied the updated model to HIV-1 using an HLA-aware framework to show how CTL response strength influences time to immune escape, as well as sequence diversity and divergence.

### Input data and simulation settings

To test our model, we simulated within-host HIV-1 evolution of *pol* and *gp120* (spliced together) using the `wavess` default values unless otherwise noted. *pol*, the polymerase gene, is under strong CTL immune pressure but not strong antibody pressure, while *gp120*, an envelope protein, is heavily targeted by the host antibody response. We used as the founder sequence *pol* and *gp120* from subtype B sequence DEMB11US006 (GenBank accession number KC473833) (Sanchez et al., 2014). We set the baseline recombination rate within each gene to *r* = 1.5 *×* 10^−5^ per breakpoint per generation, as is standard for HIV-1 (Sambaturu et al., 2025), and the rate between the two genes to *r×n*, where *n* = 1128, the number of nucleotides between *pol* and *gp120* in the founder, yielding a rate of 0.17. Conserved sites were identified as recommended in `wavess` for *pol* and *gp120* and mapped to the founder sequence. The LANL HIV database consensus sequence (Linchangco et al., 2022) was used as the reference for both genes. B-cell epitope locations were identified for *gp120* only, using the recommended method from `wavess` with the LANL HIV database ENV sequence features. The same 10 B-cell epitopes were used for all simulations.

### CTL epitopes

We identified founder T-cell epitopes for the IEDB panel of 27 HLA-A and HLA-B alleles (at the subtype, i.e. 4-digit, level) using the IEDB T-cell prediction tool with the NetMHCpan 4.1 EL prediction method for MHC-1 Binding/Elution (Reynisson et al., 2020; Vita et al., 2025). We also calculated class I pMHC immunogenicity with 1,2,C terminals masked. We filtered these epitopes based on thresholds calibrated to empirically observed numbers of escape mutations and times to maximum T-cell response, while allowing different sequences and HLAs to have different numbers of recognized epitopes. We wanted filtering criteria that did not lead to zero epitopes, but resulted in few enough to be realistic. It is estimated that between 9-18 escape mutations become fixed in the population within six months of infection (Carlson et al., 2015), with the epitope-specific time to maximum T-cell response ranging from under two weeks to over two months (Turnbull et al., 2009). With this in mind, for each set of 2 HLA-A and 2 HLA-B alleles, we identified all epitopes of length 9 amino acids under a NetMHCpan percentile of 0.1 and with an immunogenicity score of *<* 1*/*90. Across all HLA combinations, this resulted in a median of 12 epitopes across *pol* and *gp120* (range 2-25; Figure 1A). The number of days to reach maximum CTL immune cost was considered to be the inverse of the epitope immunogenicity score. Across all epitopes, the median time was 7 days (range 2-42; Figure 1B). For each epitope, the anchor positions (2 and 9) were considered positions at which a mutation would lead to escape from the CTL response (Carlson et al., 2015).

**Figure 1.**
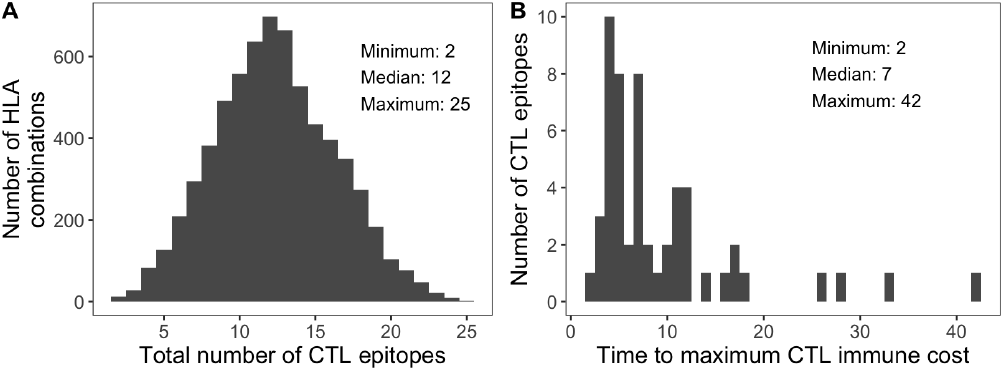
CTL response input summary. (A) Number of CTL epitopes across all combinations of two HLA-A and HLA-B molecules from the IEDB 27 CTL panel. (B) Distribution of time to maximum CTL immune response for each unique epitope from panel A.

### Simulations

For each combination of 2 HLA-A and 2 HLA-B epitopes (*n* = 6600), we simulated within-host evolution using `wavess` for one year. Weekly, we sampled 20 sequences and recorded the mean CTL fitness. For the HLA combination with the smallest number of epitopes (A*23:01, A*01:01, B*40:01, B*08:01) and one with a median number of epitopes (A*02:01, A*01:01, B*15:01, B*57:01) we also performed 100x replicates to investigate the stochasticity of CTL fitness over time. As complete immune escape may or may not occur at conserved sites for HIV-1 (Li et al., 2007; Peyerl et al., 2004), we set the conserved cost and the maximum T-cell immune cost equal to each other at a value of 0.5. To test the impact of this assumption, we also performed a sensitivity analysis on the maximum T-cell immune cost using values of 0.1, 0.3, 0.5, 0.7, and 0.9.

## Results

Overall virus fitness was strongly influenced by CTL fitness at early timepoints, with an increasing influence from other fitness factors as the viruses escaped the CTL response (Figure 2A). The overall fitness therefore started relatively low, decreased slightly at the beginning as the CTL response matured, then increased as the virus population began to escape the CTL response until it reached a relatively steady state after complete CTL escape. This results in a stark difference in early virus fitness compared to when CTL fitness is not modeled, in which case virus fitness starts off very high and decreases with increasing antibody immune pressure (Sambaturu et al., 2025).

**Figure 2.**
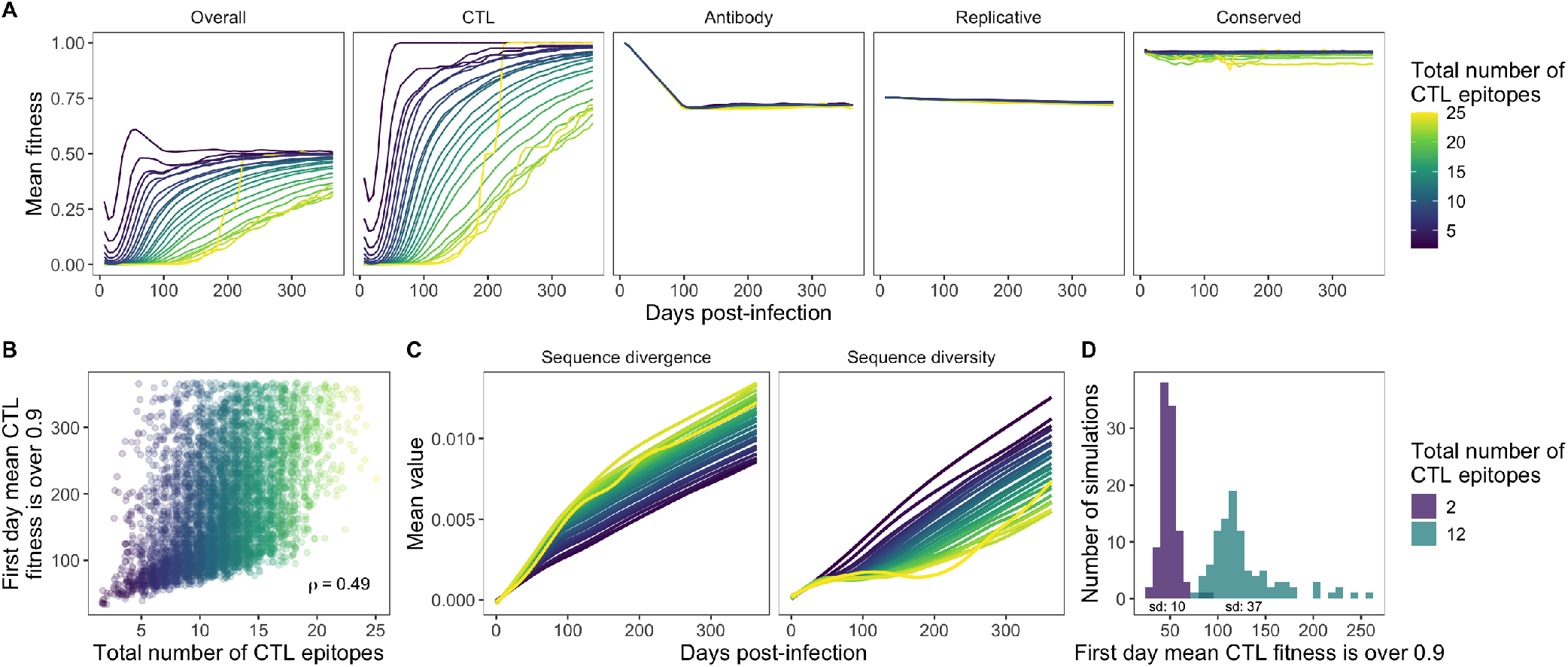
Escape from the CTL response. (A) Mean fitness across time by total number of CTL epitopes for overall fitness and each type of fitness modeled. Lines are LOESS curves with a span of 0.75. (B) Correlation between the number of CTL epitopes and the first day the mean CTL fitness was *>* 0.9. *ρ* is Spearman’s. (C) Mean sequence divergence and diversity across time by total number of CTL epitopes. Lines are LOESS curves with a span of 0.75. (D) First day the mean virus CTL fitness was *>* 0.9 for each of 100 replicates of two HLA combinations with different numbers of epitopes. The standard deviation is indicated under each histogram.

Consistent with reports of escape in real people occurring between a few weeks to several years post-infection (Goonetilleke et al., 2009), it took a median of 161 days for the mean CTL fitness to surpass 0.9 (range 35 to ≥ 365). This value was positively correlated with the number of epitopes recognized by the CTL response (Spearman *ρ* = 0.49; Figure 2B), which aligns with previous observations that HLA alleles that exert immune pressure on more sites across the HIV genome are more protective than those with less breadth (Carlson et al., 2015). Furthermore, compared to HLA combinations that recognized fewer epitopes, those that recognized more epitopes had higher sequence divergence and lower sequence diversity (Figure 2C). Finally, the time to escape for viruses with fewer recognized epitopes was less variable than the time to escape for viruses with more recognized epitopes (standard deviation of 10 days for 2 epitopes vs. 37 days for 12 epitopes; Figure 2D).

Model outputs were sensitive to the maximum CTL immune cost. First, the weaker the CTL immune response, the longer the time to escape, with higher costs resulting in stronger correlations between the total number of CTL epitopes and escape time (Figure 3A). Sequence divergence was slightly higher with increasing immune cost and sequence diversity increased more rapidly early in infection for lower costs than for higher costs, but later in infection diversity became higher on average for simulations with higher costs (Figure 3B).

**Figure 3.**
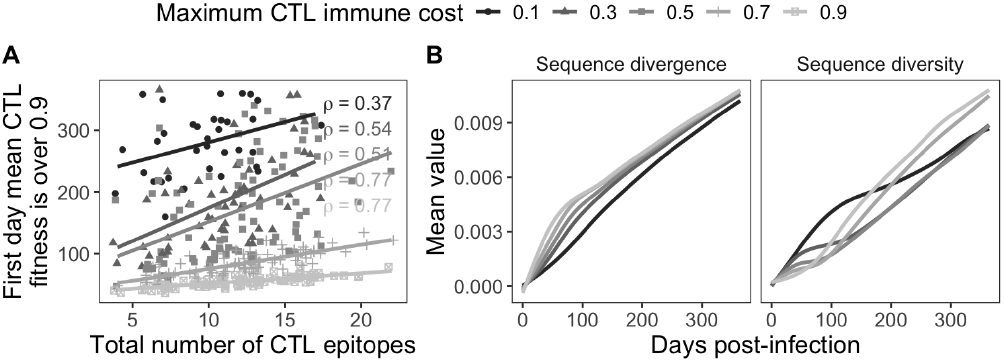
Maximum CTL immune cost sensitivity analysis. (A) Correlation between the number of CTL epitopes and the first day the mean CTL fitness was *>* 0.9. *ρ* is Spearman’s. (B) Mean sequence divergence and diversity. Lines are LOESS curves with a span of 0.75.

## Discussion

The addition of a CTL response and a variable recombination rate to `wavess` make it much more versatile than it was before. First, the framework presented here for using as model input CTL epitopes identified by the IEDB T-cell prediction tool means that the CTL response to any intracellular pathogen can be modeled, as long as a representative sequence can be obtained. Note, however, that if a virus other than HIV-1 is being modeled, the model parameters and perhaps the CTL epitope identification thresholds may have to be modified and calibrated based on empirical data for the specific pathogen of interest. Second, the addition of a variable recombination rate means that not only can recombination hotspots be included, but perhaps more importantly, non-adjacent genes or segmented sequences can also now be modeled without inaccurate hitchhiking effects. This expands the scope of `wavess` substantially, because now scenarios that can be modeled include, for example, multiple non-adjacent genes targeted by the immune system, or segmented viruses such as influenza. One limitation of the CTL response in the model is that it does not capture epitope-specific variability in immune recognition or partial immune escape. Because of this, complete CTL escape might take more time in reality, and signatures of escape in the simulated sequences may be clearer than in empirical sequences. Nevertheless, the new features added here allow for more accurate within-host modeling of virus genes under CTL-mediated selective pressures. These updates may also prove useful for modeling non-virus pathogens.

## Data availability

The data and code in this article are available at https://github.com/MolEvolEpid/wavess_ctl_manuscript.

## Author contributions statement

Z.L. and T.L. conceptualized the project, acquired funding, developed the methodology, and reviewed and edited the manuscript. T.L. supervised the project. Z.L. performed formal analysis and investigation, developed the software, visualized the results, and wrote the original draft of the manuscript.

## Acknowledgments

We thank Narmada Sambaturu, Jennifer Mamrosh, Ruy Ribeiro, and Alan Perelson for discussion about the CTL response.

